# Apparent diet digestibility of captive colobines in relation to stomach types with special reference to fibre digestion

**DOI:** 10.1101/2021.03.02.433677

**Authors:** Satoru Hoshino, Satoru Seino, Takashi Funahashi, Tomonori Hoshino, Marcus Clauss, Ikki Mastuda, Masato Yayota

**Author notes:** Corresponding author (MY).

## Abstract

Colobine monkeys are known for the anatomical complexity of their stomachs, making them distinct within the primate order. Amongst foregut fermenters, they appear peculiar because of the occurrence of two different stomach types, having either three (‘tripartite’) or four (‘quadripartite’, adding the praesaccus) chambers. The functional differences between tri and quadripartite stomachs largely remain to be explained. In this study, we aim to compare the apparent digestibility (aD) in tripartite and quadripartite colobines. Hence, we measured the aD in two colobine species, *Nasalis larvatus* (quadripartite) and *Trachypithecus cristatus* (tripartite), in two zoos. We also included existing colobine literature data on the aD and analysed whether the aD of fibre components is different between the stomach types to test the hypothesis of whether quadripartite colobines show higher aD of fibre components than tripartite colobines did. Our captive *N. larvatus* specimen had a more distinctively varying nutrient intake across seasons with a larger seasonal variation in aD than that of a pair of *T. cristatus*, which mostly consumed commercial foods with a lower proportion of browse and less seasonal variation. We observed higher aD of dry matter (DM), neutral detergent fibre (NDF) and acid detergent fibre (ADF) in the *N. larvatus* specimen, suggesting a higher gut capacity of *N. larvatus* provided by the additional praesaccus forestomach chamber. Based on the analysis of literature data for aD, we also found that quadripartite species achieved higher fibre digestibility at similar dietary fibre levels compared with tripartite species, supporting the hypothesis that the additional gut capacity offered by the praesaccus facilitates a longer retention and hence more thorough microbial fermentation of plant fibre.

## Introduction

Primates display a diverse array of digestive adaptations while covering various trophic niches, from folivory to frugivory, gummivory, insectivory, omnivory and nearly exclusive carnivory in some human populations [1, 2]. In particular, Old World monkeys of the subfamily Colobinae are known for the anatomical complexity of their stomachs, making them distinct within the primate order as the only ‘foregut fermenters’. Their stomachs are complex and multi-chambered, harbouring a symbiotic microbiome that digest plant fibre and detoxify defensive plant chemicals, possibly allowing them to exploit a diet of leaves in greater quantities than other simple-stomached primates [3, 4]. Because of such anatomical complexity with their folivory, colobines have historically often been considered difficult to maintain healthy under zoo feeding regimens, especially when compared with frugivorous and/or omnivorous primates with simple stomachs [5, 6].

In previous studies, two different types of colobine forestomach have been distinguished. The so-called ‘tripartite’ type comprises a saccus, tubiform and glandular stomach part; these can be found in all colobines. The ‘quadripartite’ type has an additional blind sac, or pouch, named ‘praesaccus’, which is thought to represent an additional chamber. It is found in the genera *Procolobus*, *Piliocolobus*, *Rhinopithecus*, *Pygathrix* and *Nasalis* [7–9]. Notably, genera with a quadripartite stomach are notoriously difficult to maintain and breed in captivity, especially in temperate regions [10–13] where constant supply of fresh browse is especially difficult in autumn and winter, and chemical composition of fresh browse differs across plant species and seasons [14]. Therefore, to reduce gastrointestinal disorders and enhance health and survival in captive colobines, identifying an appropriate diet considering nutritional differences across plant species and seasons, in relation to their digestive physiology, is one of the goals for *ex situ* animal management.

The functional differences between tri and quadripartite stomachs, however, largely remain unexplained. Matsuda, Chapman and Clauss [15] compiled literature data on the natural diet of colobine species to investigate the role of the praesaccus, suggesting that a larger gut capacity provided by an additional praesaccus is an important characteristic by which colobines survive on diets with a particularly high proportion of leaves. Thus, the higher intake capacity for species with quadripartite stomach would be assumed to be detrimental in the case of more digestible (commercial) diets in captivity than those in the wild, thereby leading to malfermentation by highly digestible components such as sugars or starch [5, 6]. Conversely, it may be assumed that species with tripartite stomachs are less susceptible to extreme bouts of malfermentation when fed highly digestible diets, simply because of their relatively reduced intake capacity. Evidently, these speculations remain to be tested.

One approach is to compare the apparent digestibility (aD), i.e. the ratio of the difference of the ingested and faecal nutrients to the ingested nutrients, in tripartite and quadripartite colobines to obtain information relevant for evaluating the digestive capacity for fibre. The aD evaluates the ability to break down and absorb nutrients, like fibre contained in browse/leaves, and thus measuring aD facilitates insights into the digestive adaptations and capacities of a species, and – maybe more importantly – also comparisons between species or species groups. It has previously been quantified in some colobines in comparison to simple-stomached primates, indicating that colobines show higher aD of fibre components, e.g. neutral detergent fibre (NDF) and acid detergent fibre (ADF), than simple-stomached primates, such as *Macaca fuscata* [16], *Alouatta* spp. [17] and *Nomascus siki* [18]. However, to our knowledge, a study focusing on comparing aD between tripartite and quadripartite colobines has not yet been undertaken. In our comparison of aD between the colobines with different stomach types, we expected that quadripartite species with a putatively larger gut capacity would display higher aD of fibre than tripartite ones.

As a first preliminary approach to this question, we examined the aD of two captive colobine species, *Nasalis larvatus* (quadripartite) and *Trachypithecus cristatus* (tripartite), in two temperate region zoos. We focused on the seasonal difference of their aD to evaluate the effects of seasonal variation in nutrient composition throughout the year on their digestive efficiency. Additionally, to test the hypothesis of whether quadripartite colobines show higher aD of fibre components than tripartite colobines, we included existing colobine literature data on the aD and analysed whether the aD of fibre components is different between the stomach types.

## Materials and methods

### Ethics statement

We conducted the feeding experiments of proboscis monkey (*N. larvatus*) and silvered langur (*T. cristatus*) in the Yokohama Zoological Gardens, Zoorasia (approval ID: #256) and Japan Monkey Centre in Japan (approval ID: #2018-016), respectively. Invasive or stressful approaches such as capture, manual restraint or anaesthesia were not performed in this study. The materials were collected from animals non-invasively. This study was approved by the Welfare of Gifu University (approval ID: #17092). All animal experiment procedures were conducted following the Guidelines for Proper Conduct of Animal Experiments (Science Council of Japan, 2006; http://www.scj.go.jp/ja/info/kohyo/pdf/kohyo-20-k16-2e.pdf) and the Guidelines of Animal Research and Welfare of Gifu University (2008; https://www.gifu-u.ac.jp/20150821-12a-experi.pdf).

### Digestion trials

In the Yokohama Zoological Gardens, the experiments were performed with one male *N. larvatus* (14 years old) housed individually. Three digestive trials were conducted in different seasons: autumn (3rd–16th September 2017), winter (2nd–15th January 2018) and summer (8th–15th June 2018). Each trial was composed of two different continuous periods: the acclimatisation (seven days) and sampling (seven days) periods. We were compelled to shorten one acclimatisation (8–12 June 2018) and sampling (13–15 June 2018) period due to heavy rain that soaked animal faeces and leftover leaves in the cage. Note that since the mean retention time for different markers in the whole digestive tract of *N. larvatus* has been reported as approximately 40 h [19], we believe that the shorter sampling period (72 h), though not ideal, is still suitable for assessing digestibility in this species. The animal was fed a mixed diet of seasonally available fresh leaves with branches and twigs harvested nearby the zoo and supplied by several commercial company or private farms (Ogawana farm, Dairi-en and Shiramori-en, Yokohama and Ohnishi Agricultural Corp. Yagazishima farm, Nago), vegetables and commercial pellets (S1 Table) two times daily. Water was freely available at all times.

In the Japan Monkey Centre, the trials were performed with two adult male *T. cristatus* (16 and 17 years old) housed together. Two experiments were conducted in different seasons: summer (30 July to 12 August 2018) and winter (14–27 February 2019) applying the same design as in *N. larvatus*, i.e. two continuous periods. Note that as their diets commonly fed did not differ in composition as much throughout the year compared to those in *N. larvatus*, we assessed these the *T. cristatus* and their diets only in two different seasons, selected for a maximum contrast of climatic conditions. The *T. cristatus* were fed a mixed diet of fresh leaves, vegetables and commercial pellets (S2 Table) three times daily. The amounts of each feed were the same between trials across the seasons. Tree leaves harvested by the zoo staffs inside the zoological garden, were fed with branch and twigs once a day at noon, and the other feeds provided by commercial suppliers, were fed in the morning and evening.

We measured the body mass of all animals before and after the sampling periods in each experiment (*N. larvatus*: Digital Platform Scale, DP-8100, Yamato, Japan; *T. cristatus*: SD75LJP, OHAUS Corporation, USA), by offering weighing platforms on which the animals stepped voluntarily.

### Sampling procedures

Feed intake was quantitatively recorded over seven sampling days. Each food item was weighed before it was offered to the animals and left in their enclosures until the next feeding session. S1-S2 Table shows the mean (±standard error) daily amount of offered food per animal. All leftover food was removed, and the enclosure was cleaned before fresh food items were offered. All food items and leftovers were weighed with accuracy of 2 g (browse, UDS-500N, Yamato, Japan) or 1 g (others, UH-3201, A&D Company, Japan). Leftover weights were adjusted by deriving a desiccation factor from the measured moisture lost from similar sets of food placed in a desiccation pan in an area adjacent to the primate enclosures. For the two *T. cristatus* at Japan Monkey Centre housed together, individual feed intake and faeces output were calculated as the average of total measures divided by two.

We collected equal amounts of each feedstuff at every feeding time during the sampling period and stored them in a refrigerator (10°C: trials for *N. larvatus*; 4°C: trials for *T. cristatus*) and a freezer (−18 to 20°C: only leaves and banana peel for *T. cristatus*) for nutritional analysis. We also collected all faeces shortly before every feeding time and immediately preserved them in the freezer. We mixed each feed sample and then collected 100 g of leaves of each browse species and all amounts of other foods as representative samples. We mixed all faeces with 1 ml/100 g of 10% formalin solution and preserved 500 g faeces as a representative sample [20].

After measuring the fresh weight of representative samples, in the case of *N. larvatus*, we lyophilised leaf and faeces samples using a freeze-dryer (DC400/800, Yamato, Japan). For other feedstuff samples (vegetables and commercial pellets), we used an air-forced dry oven (DKM812, Yamato, Japan) for 48 h at 60°C. Note that we mashed soybean and peanuts in a mortar due to their high fat content and then soaked and washed the mixture with ether to extract fat. After fat extraction, the soybean and peanuts were dried using an air-forced dry oven (DKM600, Yamato, Japan) for 3 h at 60°C. In the case of *T. cristatus*, we dried all feedstuff and faecal samples using an air-forced dry oven (DKM812, Yamato, Japan) for 48 h at 60°C.

We ground leaf and faecal samples using a Wiley mill through a 2-mm screen and ground other feed samples using a coffee mill to avoid heat denaturation of sugars in the feeds.

### Estimation of apparent digestibility (aD)

We used the average daily feed intake and faecal output during the sampling term to estimate the aD in each trial. We analysed the dry matter (DM), crude ash (CA), crude protein (CP) and acid detergent fibre (ADFom) in accordance with AOAC 930.15, 942.05, 990.03 and 973.18, respectively [21]. We then analysed neutral detergent fibre (aNDFom) according to the method of van Soest, Robertson and Lewis [22]. Detergent fibre data are presented without residual ash. We calculated aD of each nutrient (N) according to the following equation: aD_N_ (%) = (N_feed intake_–N_feces_) / N_food intake_ × 100.

### Literature data and analysis

Together with our new experimental data, as shown in Table 1, we used published aD data of seven colobine species, including four tripartite species (13 datasets) [16, 17, 23–26] and two quadripartite species (five datasets) [17, 18, 27] to compare digestive capacity between the stomach groups. These published data included total feed and fibre intake and aD of DM, NDF and ADF, and for some but not all studies the body mass of the animals used.

**Table 1.**
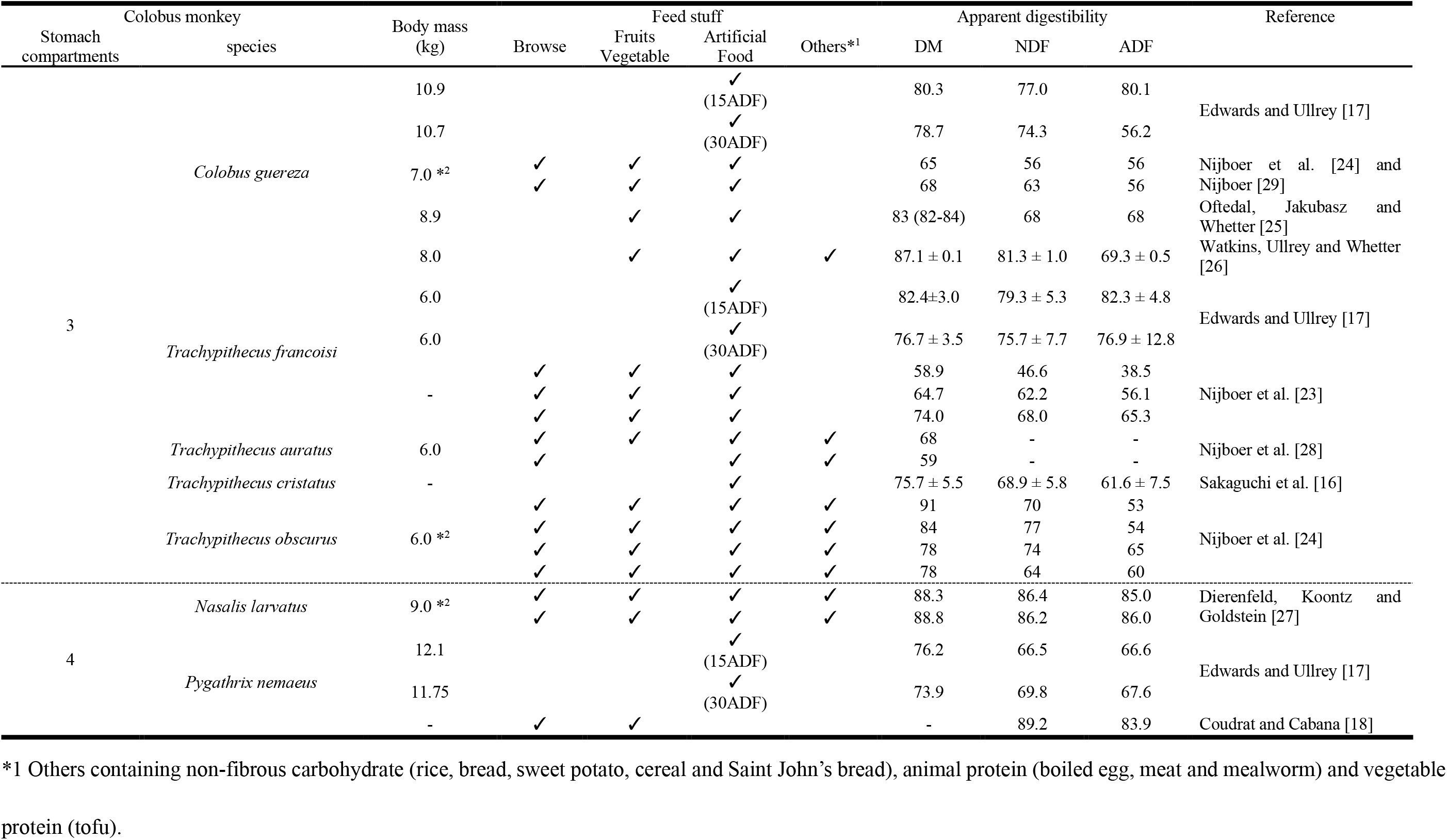
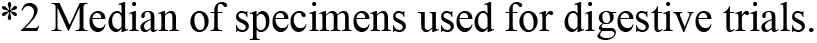
Summary of colobine apparent digestibility and body mass in the previous studies.

Due to the limited data, we only focussed on two-factorial models and did not perform models with more variables/factors. First, we tested two-factorial linear models that linked the aD to either combination of body mass, stomach type and NDF content of the ingested diet. Second, in the larger dataset (including the studies did not provide body mass information), we examined the relationship between aD in DM, NDF and ADF and the dietary NDF content in each colobine group with different stomach type to estimate the slope and intercept using linear regression analysis. Analysis of covariance served to compare the slopes and intercepts of regression lines between the stomach types. Note that we could not conduct the same statistical analysis for the aD of CP because of the lack of published data. Additionally, because Nijboer et al. [28] analysed only crude fibre as fibre contents and Coudrat and Cabana [18] did not specify DM contents in diets and faeces with the total intake (see Table 1), we could not include these data in our analysis. All statistical analyses were performed in Spyder (Python 3.7).

## Results

### Digestion trials

The body mass of *Nasalis larvatus* in autumn, winter and summer was 17.3, 18.9 and 17.8 kg, respectively, shortly before beginning of the sampling periods. The composition and amount of the diet were different amongst trials (S1 Table). The ratio of leaves/other feed intake were 7.4:1, 3.3:1 and 3.5:1, in autumn, winter and summer. Browse species generally contained more fibre than other feeds (S3–S5 Table), possibly leading to different proportion of ADFom in the diet in autumn, i.e. 33.3%, 29.4% and 30.1% of DM in autumn, winter and summer, respectively. There was notable seasonal variation in the nutritional composition of the browse species (S3–S5 Table). The DM and ash contents of laurel (*Machilus thunbergii*) in winter were 5%–10% higher than those in summer and winter. The aNDFom and ADFom of each browse species in winter were 5%–10% lower than those in summer and autumn. On the other hand, nutrient contents of fruits, vegetables, beans, starchy foods and commercial products in *N. larvatus* did not differ between the seasons (S3–S5 Table). The aD of DM was 69.9%, 79.6% and 73.7% in autumn, winter and summer, respectively (Table 2). Likewise, the aD of CP, aNDFom and ADFom varied amongst the three seasons (Table 2).

**Table 2.**
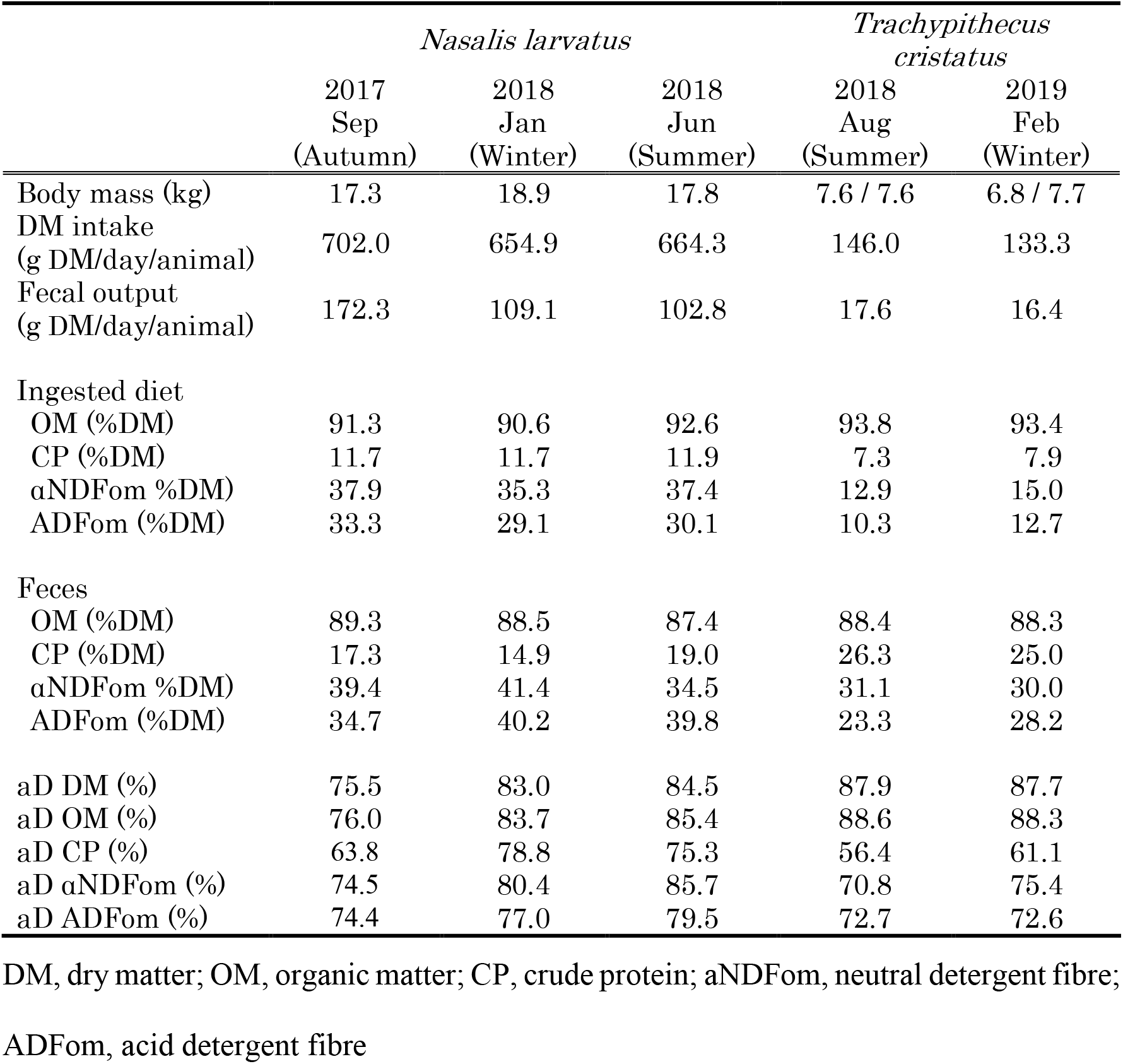
DM intake, faecal output and apparent digestibility (aD) in *Nasalis larvatus* and *Trachypithecus cristatus*.

The body mass of the two *T. cristatus* was 7.6 and 7.6 kg in summer and 6.8 and 7.7 kg in winter shortly before beginning of the sampling period. There was strong seasonality in the nutritional composition of their browse (S6–S7 Table). The DM and ash contents in the browse species in *T. cristatus* were higher, and CP, aNDFom and ADFom were lower in winter than in summer. The other diet items, i.e. fruits, vegetables, starchy foods and commercial product, did not differ between the two seasons (S6–S7 Table). The aD of DM, ash and ADFom did not differ between the two seasons; only the aD of CP and aNDFom changed between the two seasons (Table 2).

### Literature data

#### Apparent digestibility

Body mass differed significantly between species with a tripartite (mean = 7.39 kg; sd = 0.71) or a quadripartite (13.7 kg; sd = 4.23) stomach (Z = 3.40, p < 0.001).

In a model linking aD DM with body mass and stomach type (R^2^ = 0.016, p = 0.862), neither body mass (F=0.015, p = 0.903) nor stomach type (F= 0.072, p= 0.792) were significant. However, for aD NDF and aD ADF, particularly stomach type approached significance in both models, and body mass in the model for aD ADF (whole model: R^2^ = 0.170, p = 0.187 and R^2^ = 0.252, p = 0.074; body mass F = 2.252, p = 0.150 and F = 3.025, p = 0.074; stomach type: F = 3.866, p = 0.064 and F = 3.025, p = 0.074, respectively).

On the other hand, in a model linking aD DM with body mass and NDF contents of the ingested diet (R^2^ = 0.571, p = 0.0005), only the NDF content (F= 22.04, p < 0.001) was significant but not body mass (F = 0.015, p = 0.903). For aD NDF and aD ADF, the neither body mass nor NDF content were significant (whole model: R^2^ = 0.287, p = 0.047 and R^2^ = 0.364, p = 0.017; body mass: F = 2.252, p = 0.150 and F = 2.998, p = 0.100; NDF contents: F = 2.138, p = 0.160 and F = 2.744, p = 0.114, respectively). Note that in comparison to the models presented further below, sample sizes were limited as not all literature provided data on body mass.

The final model for the limited dataset evaluated the effects of aD DM on the stomach type and NDF content (R^2^ = 0.550, p = 0.001), where only the NDF content (F= 22.04, p = 0.0002) was significant but not stomach type (F = 0.072, p = 0.792). For aD NDF and aD ADF, only stomach type was significant or approached significance (whole model: R^2^ = 0.291, p = 0.045 and R^2^ = 0.406, p = 0.009; stomach type: F = 3.866, p = 0.064 and F = 6.382, p = 0.021; NDF contents: F = 2.138, p = 0.160 and F = 2.744, p = 0.114, respectively).

In the larger dataset, there was a significant negative relationship between the aD of DM and the NDF content of the ingested diet of six colobine species (r = −0.770, p < 0.001). The difference in the regression slopes and intercepts for aD of DM vs. NDF content of the ingested diet between the stomach types was not significant (Fig 1; slope, t = 1.079; p = 0.293; intercept, t = 1.092, p = 0.287). There was also a significant negative relationship between the aD of NDF or the aD of ADF and the NDF content of the ingested diet of six colobine species (NDF, r = −0.459, p = 0.021; ADF, r = −0.439, p = 0.028). The differences in the regression intercepts for both aD of NDF and aD of ADF vs. NDF content were significant (NDF, t = −2.559, p = 0.018; ADF, t = −2.487, p = 0.021) between the stomach types (Fig 1), while the slopes were not different (NDF, t = 0.267, p = 0.792; ADF, t = 0.152, p = 0.881).

**Fig 1.**
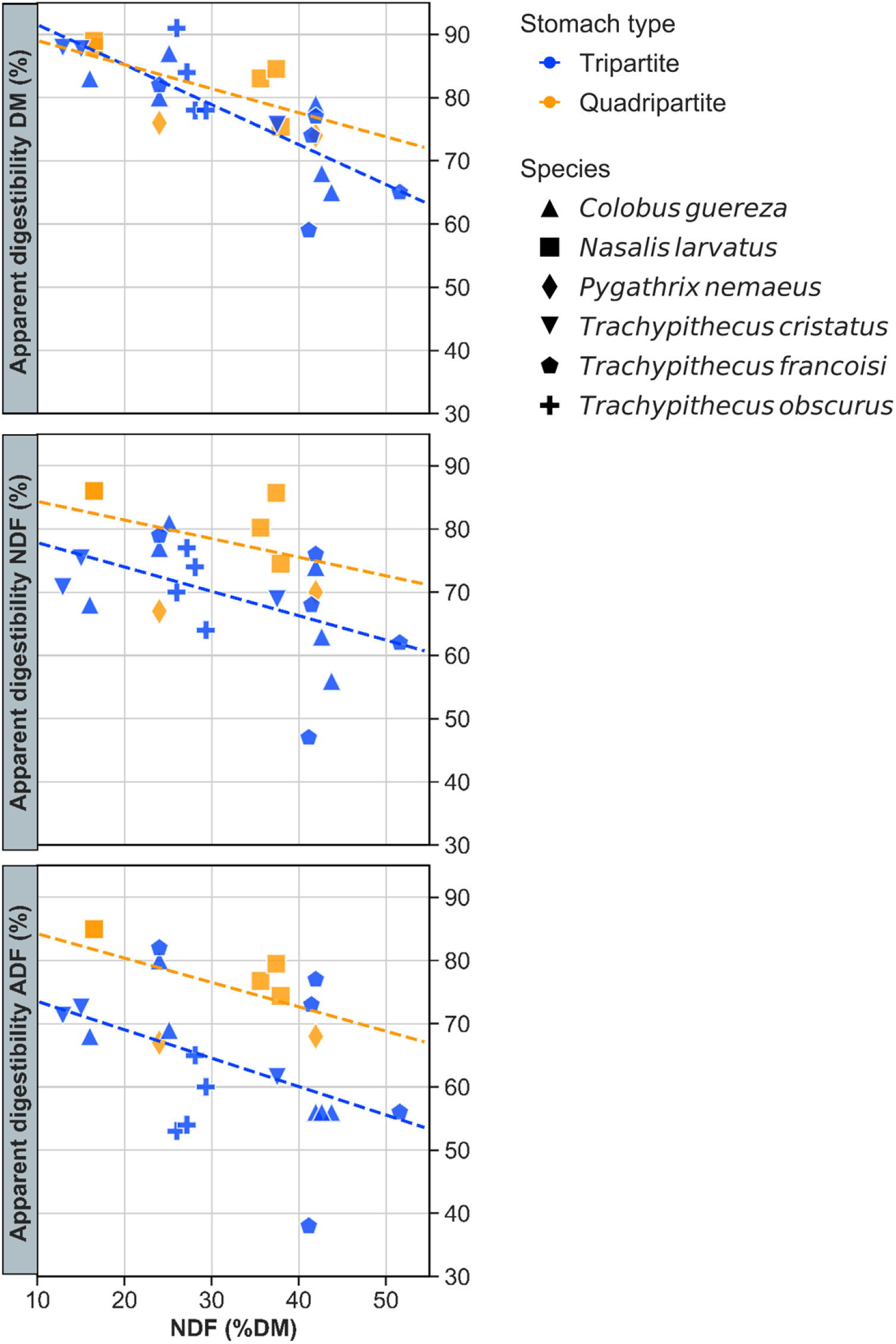
Relationship between apparent digestibility of dry matter (DM, %), neutral detergent fibre (NDF, %) or acid detergent fibre (ADF, %) and neutral detergent fibre intake (%) in six colobine species: *Colobus guereza* [17], *Nasalis larvatus* [27], *Pygathrix nemaeus* [17], *Trachypithecus cristatus* [16], *Trachypithecus francoisi* [17, 23] and *Trachypithecus obscurus* [24]. Species and stomach type, i.e. tri- or quadripartite, are indicated in different shape and colour symbols, respectively.

## Discussion

As expected, we confirmed the variation in the aD because of the nutrient differences in the diets across the seasons. Each browse species’ nutrient composition changed seasonally, as reported for deciduous tree species in North American zoos [14], whereas those of commercial food items were stable (S3–S7 Table). The aD of DM, aNDFom and ADFom in *N. larvatus* clearly changed across seasons: aD DM by 9.1%, aD aNDFom by 11.2 % and aD ADFom by 5.1%. On the other hand, aD was relatively stable between the two different seasons in *T. cristatus*, although the total DM intake in winter was slightly higher than that in summer. As in the present study, Edwards and Ullrey [17] noted that the aD of DM in three colobine species (*Colobus guereza*, *Pygathrix nemaeus* and *Trachypithecus francoisi*) decreased with an increased fibre level in the diet, supporting that aD is affected by a variation in food composition and seasonally varying nutrient contents in feeds. Consequently, the aD of *N. larvatus* had a larger seasonal variation, with more distinctively varying nutrient intake across seasons, than that of *T. cristatus*, which mostly consumed commercial foods with a lower proportion of browse. Evaluating the nutrient digestibility of captive colobines on different diets throughout the year may contribute to their health management and predict intake requirements across different diets and seasons.

Comparison of the nutrient composition (especially fibre) of faeces in free-ranging and captive individuals has been proposed to obtain information relevant for the improvement of diets of colobines [10]. *Nasalis larvatus* in our study had faecal NDF contents (35%–41% in DM) that were higher than those reported in other captive conspecifics, i.e. 17% [27] (mean of two different values), but lower than those of free-ranging ones, i.e. 53%–70% [10]. Although faecal NDF contents of free-ranging *T. cristatus* are not available, those in our study (30%–31%) were comparable to other closely related species in captivity, i.e. 37% (mean of six different values in *T. auratus*) [24] and 31% (mean of three different values in *T. francoisi*) [23, 29], although still far lower than those reported for free-ranging *N. larvatus*. Altering the diets of captive colobines to include more fibre, comparable to those of free-ranging ones, may be recommendable.

In the present study, the NDF level of the total DM intake was much higher in the quadripartite species *N. larvatus* (35.6%–37.9%) than in the tripartite species *T. cristatus* (12.9%–15.0%), because of a much higher proportion of browse fed to the former. This zoo practice may stem from the impression that quadripartite species are generally more difficult to maintain in captivity; therefore, more effort is undertaken to provide them with feed items considered natural for them, mainly browse. Correspondingly, we observed a higher DM intake (% BW, shown in S3–S5 Table) in *N. larvatus* compared with those in *T. cristatus* that could be interpreted as compensation for the higher fibre levels. However, regardless of the higher fibre and intake levels, we observed higher aD in DM, aNDFom and ADFom (Table 2) in the *N. larvatus* specimen. This observation is most parsimoniously explained by a higher gut capacity in the proboscis monkey, provided by the additional praesaccus forestomach chamber. Typically, a higher relative food intake leads to shorter digesta retention times and can also compromise digestibility [30, 31, cf. also Fig 2]. However, a higher gut capacity can mitigate this effect, and this may be the main adaptive value of the praesaccus in quadripartite species [15].

**Fig 2.**
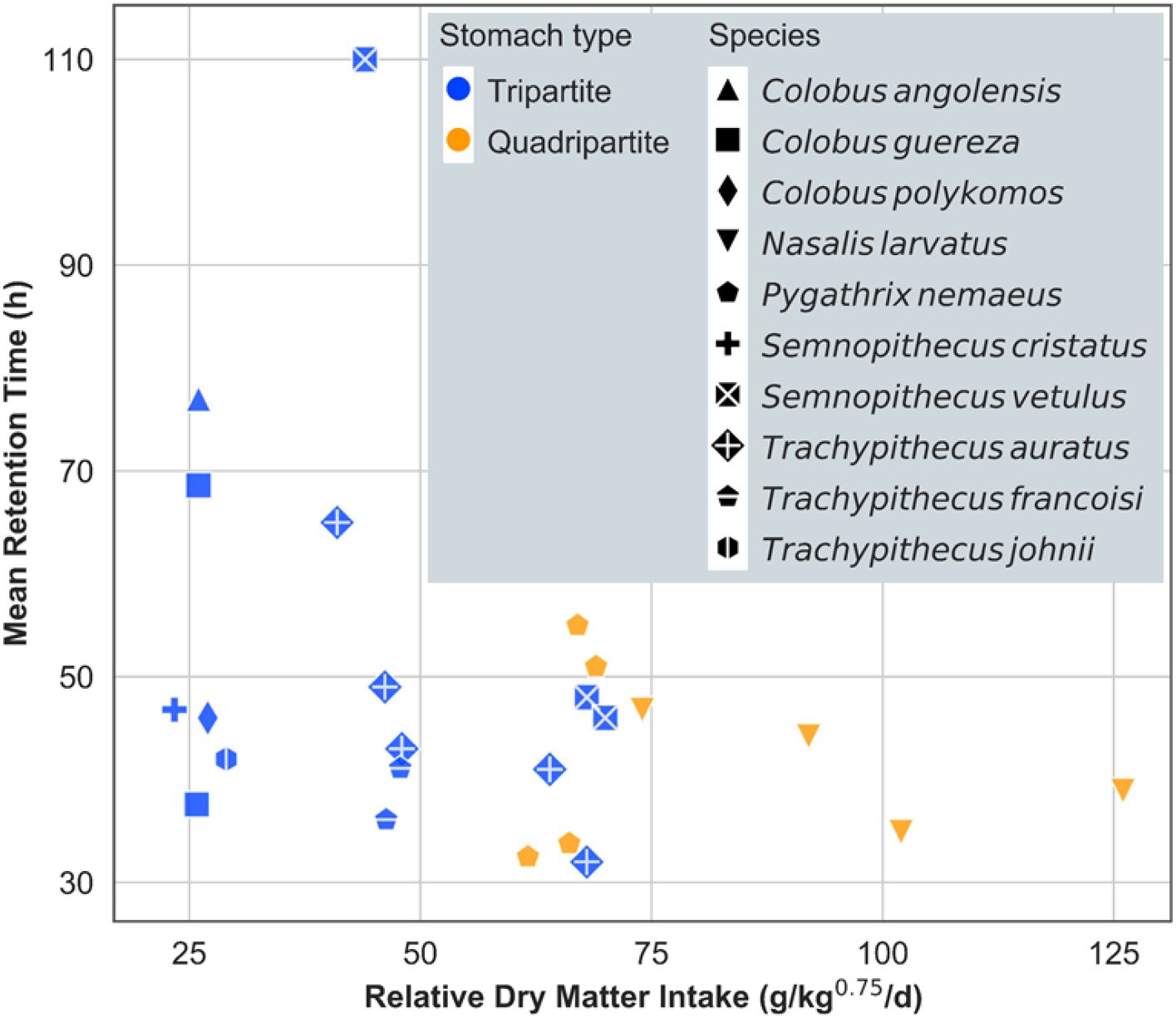
Relationship between mean retention time (h) and relative dry matter intake (g/kg^0.75^/d) in 10 colobine species: *Colobus angolensis* [47], *Colobus guereza* [17], *Nasalis larvatus* [19], *Pygathrix nemaeus* [17], *Semnopithecus cristatus* [16], *Semnopithecus vetulus* [48], *Trachypithecus auratus* [28], *Trachypithecus francoisi* [17] and *Trachypithecus johnii* [47]. Species and stomach type, i.e. tri- or quadripartite, are indicated in different shape and colour symbols, respectively.

Unfortunately, the currently available information for tri- and quadripartite species is biased in terms of the body mass of the investigated specimens. Although the overall distribution of tri- and quadripartite stomach types across colobine genera [15] does not suggest a body size effect on stomach anatomy, quadripartite specimens of the literature data collection were significantly heavier than tripartite specimens. This is due to the inclusion of *N. larvatus* – the largest colobine, with a quadripartite stomach. Given an older but prevailing assumption in the ecophysiological literature that body mass is positively related to digesta retention time and digestibility [32], this could lead to the suspicion that differences between the stomach types are body size effects. However, this is refuted on the one hand by our own analyses that showed no effect of body mass or a trend subordinate to the trend of stomach type, and on the other hand by more recent literature. An effect of body size on digesta retention and digestion has been contested by various large empirical data collections as well as by theoretical considerations [31, 33, 34], including exclusive analyses of primate data.

The results of our analysis for aD using the larger literature data set further suggested functional differences between the stomach types. There was no difference in how dietary fibre content influenced the overall DM digestibility, which is, in most of the diets used in these studies, most likely dominated by the digestion of non-fibrous carbohydrates and protein. However, stomach type had a significant effect on how dietary fibre levels affected the digestibility of fibre itself. Here, quadripartite species achieved higher fibre digestibility at similar dietary fibre levels than tripartite species, suggesting that the additional gut capacity offered by the praesaccus facilitates longer retention and hence more thorough microbial fermentation of plant fibre. One notable tendency was that *N. larvatus* achieved a particularly high digestibility, driving the difference between the two stomach types. In this species, regurgitation and remastication have been observed in the wild [35], and a smaller faecal particle size of *N. larvatus* compared with other colobines has been reported [36]. Because particle size reduction via chewing is one of several key factors affecting digestibility [37], we cannot determine whether the difference in digestibility observed here is related to stomach type or chewing efficiency. Ideally, in future studies, similar diets should be employed as this will allow meaningful comparison of faecal particle size; chewing behaviour should also be observed.

Unfortunately, the current available data on digesta retention times in colobine species does not allow testing for a general difference between tripartite and quadripartite species (Fig 2). Between species, retention times need to be compared in relation to the food intake level [30, 38]. However, the ranges of intake level in published studies hardly overlap between the stomach groups, making a reasonable comparison impossible. To test whether quadripartite species achieve longer digesta retention because of a higher gut fill, comparative studies with different species on a similar (possibly browse-dominated) diet would be required where intake, digestibility and digesta retention are assessed in the same experiment, additionally facilitating the calculation of gut fill [39, 40]. Ideally, such a study would also address the problem of the very limited sample size of the present experiments. Until such a study is performed, our results must be considered preliminary, delivering plausible hypotheses.

It should be noted that we cannot exclude the possibility of the effects of specific fibre-digesting bacteria in the presaccus in quadripartite species. So, far, only a few analyses of the forestomach microbiome are available for colobines. Although the recent developments in sequencing technology describe the foregut microbiome in some colobines, e.g. *N. larvatus* [41, 42] and *Rhinopithecus roxellana* [43], the function of these microbe species has not been evaluated. However, there is currently also no reason to assume that the praesaccus should harbour a fundamentally different microbiome from the saccus. Detailed studies about differences in the microbiome at different forestomach locations, as available in ruminants [e.g. 44], do not exist for colobines so far. In contrast to the detailed knowledge about the differential function of individual forestomach sections in ruminants and camelids [e.g. 45], there is, to date, no indication of the differential function of the forestomach compartments of non-ruminant foregut fermenters [46] beyond the provision of sheer fermentation chamber capacity, and this may well also apply to colobines.

## Acknowledgements

Our appreciation goes to the animal caretakers and veterinarians from Zoorasia and Japan Monkey Centre, namely, Ryuta Kawasaki and Kei Watanabe, without whom this project would not have been possible. General supervision and administrative supports were provided by Dr. Takashi Hayakawa and Dr. Koshiro Watanuki as the counterparts of Japan Monkey Centre. In particular, S.H. thanks to Dr. T. Hayakawa for his kind arrangement of the freezer to preserve the collected samples at his laboratory of the Primate Research Institute, Kyoto University and for his technical editing of this manuscript.

## Author contributions

SH, IM, MC and MY conceptualised the idea and drafted the manuscript; SH performed the feeding trials; SH, IM, MC and MY performed and interpreted the statistical analysis; SS, TF and TH arranged the sampling in the zoos and MY organised the projects. All authors contributed to the final version of the manuscript.

## Supporting information

**S1 Table.** Feed composition, and offered and leftover amounts (g as fed/day) in each digestion trial of an adult male of *Nasalis larvatus*.

|  |  | 2017 Sep (Autumn) |  | 2018 Jan (Winter) |  | 2018 Jun (Summer) |  |
| --- | --- | --- | --- | --- | --- | --- | --- |
|  |  | Offered*<sup>1</sup> | Leftover*<sup>1</sup> | Offered*<sup>1</sup> | Leftover*<sup>1</sup> | Offered*<sup>1</sup> | Leftover*<sup>1</sup> |
| Browse*<sup>2</sup> | Yoshino cherry (*Prunus yedoensis* Matumura) | 1551±120 | 996±128 | - | - | 1537±420 | 1114±287 |
|  | Bamboo-leaf oak (*Quercus myrsinifolia*) | 1708±221 | 1441±190 | 2111±279 | 1930±276 | - | - |
|  | Chinquapin (*Castanopsis sieboldii*) | 1611±146 | 1405±124 | 969±110 | 358±264 | - | - |
|  | Laurel (*Machilus thunbergii*) | 1077±126 | 846±99 | 2583±426 | 2113±373 | 3317±931 | 2682±619 |
|  | Glossy privet (*Ligustrum lucidum*) | 1339±130 | 1177±108 | 1040±73 | 894±77 | 1799±626 | 1637±595 |
|  | Willow (*Salix spp.*) | 761±139 | 452±96 | - | - | - | - |
|  | Hibiscus (*Hibiscus spp.*) | - | - | 187±22 | 58±8 | - | - |
|  | Japanese spindletree (*Euonymus japonicus*) | - | - | 768±35 | 571± 39 | 758±88 | 382±131 |
| Fruit | Apple | 129±6 | 0 | 163±1 | 3±2 | 167±5 | 0 |
| Vegetable | Carrot | 123±2 | 1±1 | 125±1 | 93±2 | 122±13 | 14±7 |
|  | Green bean | 124±2 | 2±1 | 173±3 | 101±5 | 158±2 | 9±2 |
|  | Broccoli | 108±2 | 0 | 148±3 | 3±2 | 122±12 | 0 |
|  | Asparagus | 104±1 | 0 | 101±1 | 4±1 | 77±8 | 2±2 |
|  | Cucumber | 199±4 | 5±5 | 121±6 | 3±3 | 97±5 | 0 |
| Beans | Soy bean | 31±1 | 0 | 41±1 | 0 | 37±2 | 0 |
|  | Peanuts | 25±1 | 6±1 | 73±2 | 26±1 | 61±4 | 29±0 |
| Commercial product | Primate L/S biscuit banana (Mazuri) | 16±1 | 2±0 | 141±2 | 2±1 | 58±0 | 0 |

\*<sup>1</sup> Weight unit is X ± SE g, as fed/day/animal.
\*<sup>2</sup> We measured the weight of the whole branch, including leaves and twigs.

**S2 Table.** Feed composition, and offered and leftover amounts (g as fed/day/two animals) in each digestion trial of two adults of *Trachypithecus cristatus*.

|  |  | 2018 Aug<br>(Summer) |  | 2019 Feb<br>(Winter) |  |
| --- | --- | --- | --- | --- | --- |
|  |  | Offered*<sup>1</sup> | Leftover*<sup>1</sup> | Offered*<sup>1</sup> | Leftover*<sup>1</sup> |
| Browse*<sup>2</sup> | Bamboo-leaf oak (*Quercus myrsinifolia*) | 300±0 | 269±5 | 300±1 | 280±4 |
| Fruit | Apple | 600±1 | 81±16 | 600±0 | 1±1 |
|  | Banana peel | 80±0 | 3±1 | 80±0 | 0 |
| Vegetable | Carrot | 60±1 | 1±1 | 60±0 | 0 |
|  | Cabbage | 900±0 | 69±17 | 901±0 | 9±2 |
| Starchy food | Sweet potato | 320±0 | 0 | 321±0 | 0 |
| Commercial product | Leaf-eater primate diet - Mini-biscuit (Mazuri) | 20±0 | 0 | 20±0 | 0 |

\*1 Weight unit is X ± SE g FM/day.
\*2 We measured the weight of the whole branch, including leaves and twigs.

**S3 Table.**
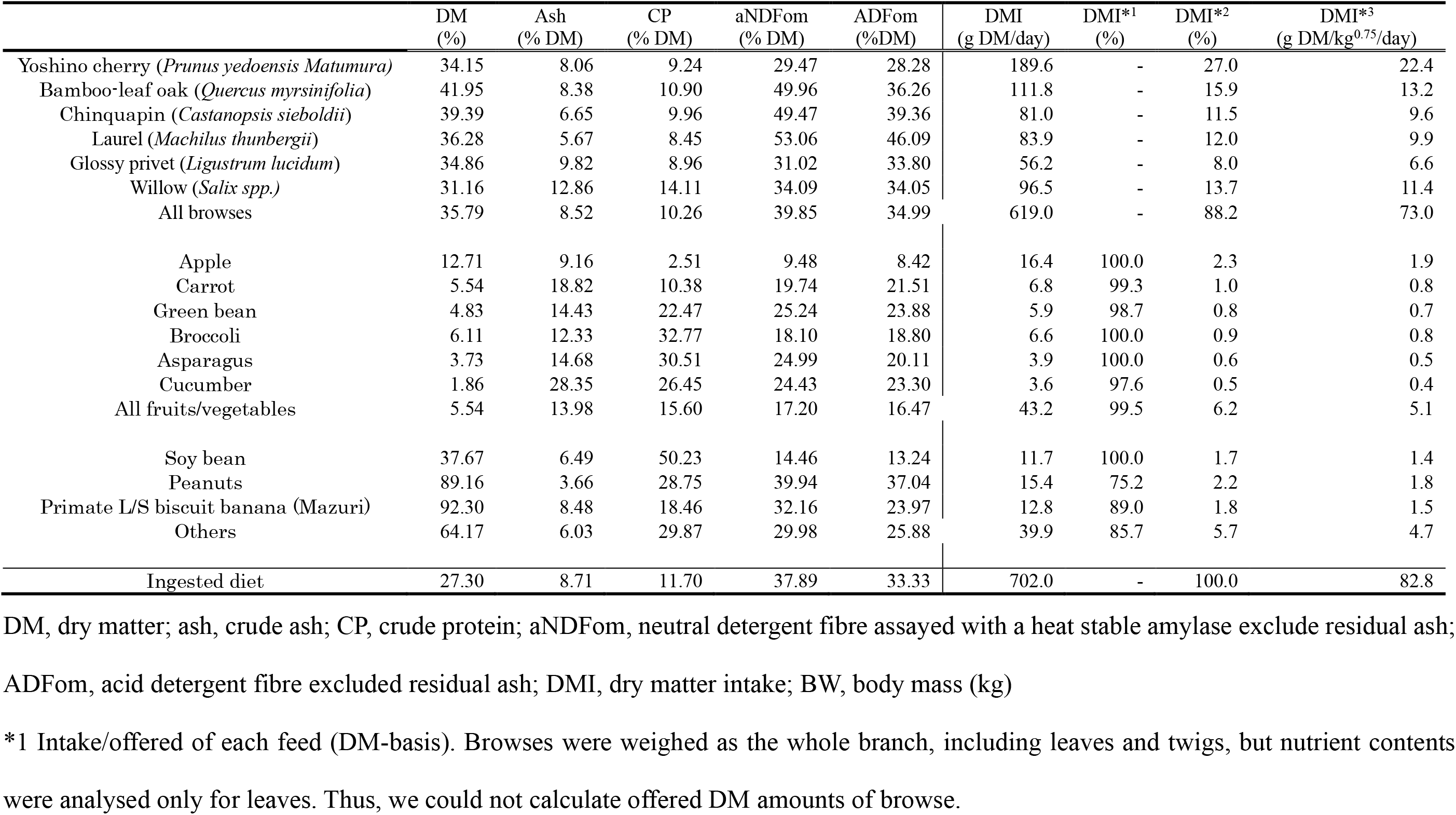
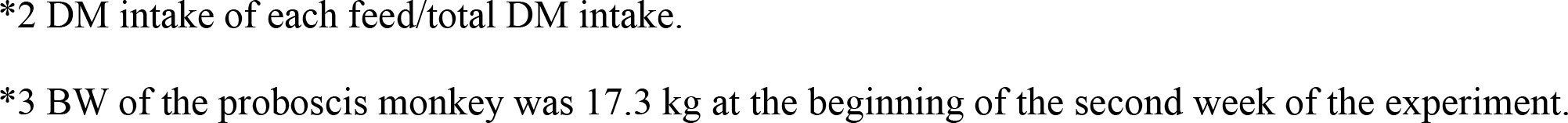
Nutrient contents in the feeds and DM intake of an adult male of *Nasalis larvatus* in September 2017 (autumn)

|  | DM<br>(%) | Ash<br>(% DM) | CP<br>(% DM) | aNDFom<br>(% DM) | ADFom<br>(%DM) | DMI<br>(g DM/day) | DMI*<sup>1</sup><br>(%) | DMI*<sup>2</sup><br>(%) | DMI*<sup>3</sup><br>(g DM/kg<sup>0.75</sup>/day) |
| --- | --- | --- | --- | --- | --- | --- | --- | --- | --- |
| Yoshino cherry (*Prunus yedoensis* Matumura) | 34.15 | 8.06 | 9.24 | 29.47 | 28.28 | 189.6 | - | 27.0 | 22.4 |
| Bamboo-leaf oak (*Quercus myrsinifolia*) | 41.95 | 8.38 | 10.90 | 49.96 | 36.26 | 111.8 | - | 15.9 | 13.2 |
| Chinquapin (*Castanopsis sieboldii*) | 39.39 | 6.65 | 9.96 | 49.47 | 39.36 | 81.0 | - | 11.5 | 9.6 |
| Laurel (*Machilus thunbergii*) | 36.28 | 5.67 | 8.45 | 53.06 | 46.09 | 83.9 | - | 12.0 | 9.9 |
| Glossy privet (*Ligustrum lucidum*) | 34.86 | 9.82 | 8.96 | 31.02 | 33.80 | 56.2 | - | 8.0 | 6.6 |
| Willow (*Salix spp.*) | 31.16 | 12.86 | 14.11 | 34.09 | 34.05 | 96.5 | - | 13.7 | 11.4 |
| All browses | 35.79 | 8.52 | 10.26 | 39.85 | 34.99 | 619.0 | - | 88.2 | 73.0 |
| Apple | 12.71 | 9.16 | 2.51 | 9.48 | 8.42 | 16.4 | 100.0 | 2.3 | 1.9 |
| Carrot | 5.54 | 18.82 | 10.38 | 19.74 | 21.51 | 6.8 | 99.3 | 1.0 | 0.8 |
| Green bean | 4.83 | 14.43 | 22.47 | 25.24 | 23.88 | 5.9 | 98.7 | 0.8 | 0.7 |
| Broccoli | 6.11 | 12.33 | 32.77 | 18.10 | 18.80 | 6.6 | 100.0 | 0.9 | 0.8 |
| Asparagus | 3.73 | 14.68 | 30.51 | 24.99 | 20.11 | 3.9 | 100.0 | 0.6 | 0.5 |
| Cucumber | 1.86 | 28.35 | 26.45 | 24.43 | 23.30 | 3.6 | 97.6 | 0.5 | 0.4 |
| All fruits/vegetables | 5.54 | 13.98 | 15.60 | 17.20 | 16.47 | 43.2 | 99.5 | 6.2 | 5.1 |
| Soy bean | 37.67 | 6.49 | 50.23 | 14.46 | 13.24 | 11.7 | 100.0 | 1.7 | 1.4 |
| Peanuts | 89.16 | 3.66 | 28.75 | 39.94 | 37.04 | 15.4 | 75.2 | 2.2 | 1.8 |
| Primate L/S biscuit banana (Mazuri) | 92.30 | 8.48 | 18.46 | 32.16 | 23.97 | 12.8 | 89.0 | 1.8 | 1.5 |
| Others | 64.17 | 6.03 | 29.87 | 29.98 | 25.88 | 39.9 | 85.7 | 5.7 | 4.7 |
| Ingested diet | 27.30 | 8.71 | 11.70 | 37.89 | 33.33 | 702.0 | - | 100.0 | 82.8 |

DM, dry matter; ash, crude ash; CP, crude protein; aNDFom, neutral detergent fibre assayed with a heat stable amylase exclude residual ash;
ADFom, acid detergent fibre excluded residual ash; DMI, dry matter intake; BW, body mass (kg)
\*1 Intake/offered of each feed (DM-basis). Browses were weighed as the whole branch, including leaves and twigs, but nutrient contents were analysed only for leaves. Thus, we could not calculate offered DM amounts of browse.
\*2 DM intake of each feed/total DM intake.
\*3 BW of the proboscis monkey was 17.3 kg at the beginning of the second week of the experiment.

**S4 Table.**
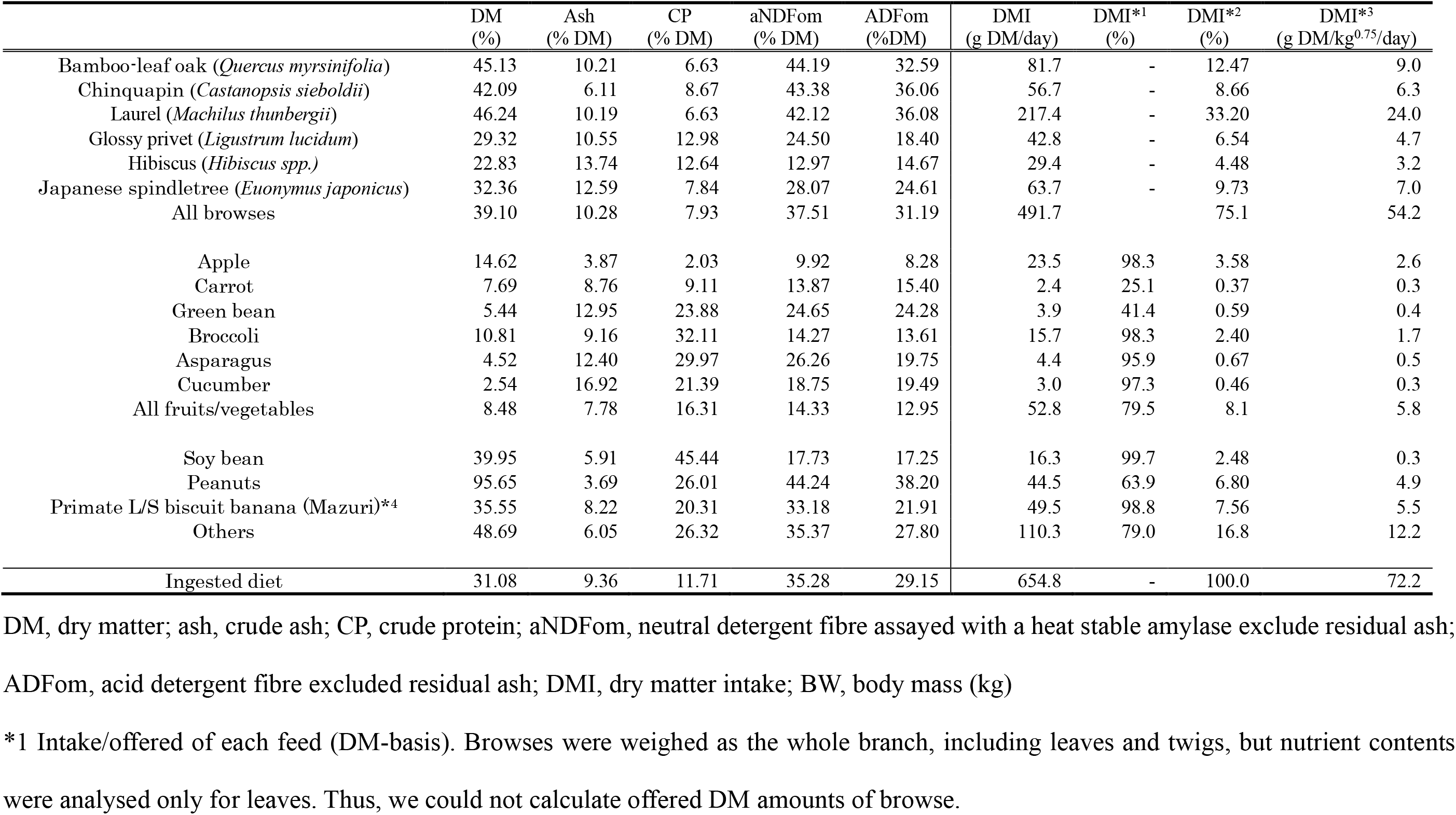
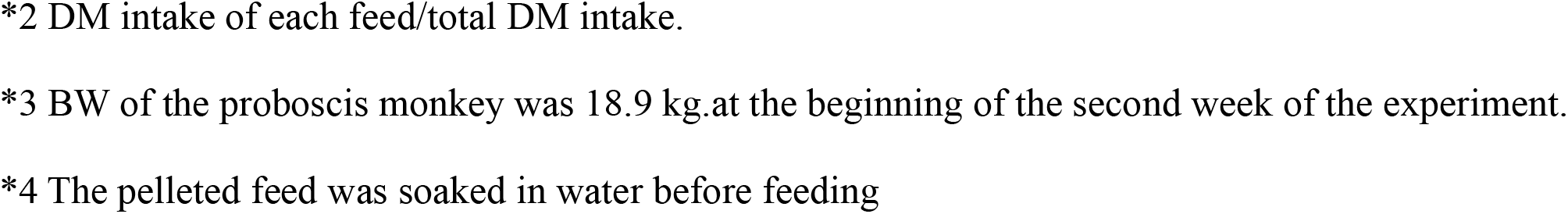
Nutrient contents in the diet and DM intake of an adult male of *Nasalis larvatus* in January 2018 (winter)

|  | DM<br>(%) | Ash<br>(% DM) | CP<br>(% DM) | aNDFom<br>(% DM) | ADFom<br>(%DM) | DMI<br>(g DM/day) | DMI*1<br>(%) | DMI*2<br>(%) | DMI*3<br>(g DM/kg<sup>0.75</sup>/day) |
| --- | --- | --- | --- | --- | --- | --- | --- | --- | --- |
| Bamboo-leaf oak (*Quercus myrsinifolia*) | 45.13 | 10.21 | 6.63 | 44.19 | 32.59 | 81.7 | - | 12.47 | 9.0 |
| Chinquapin (*Castanopsis sieboldii*) | 42.09 | 6.11 | 8.67 | 43.38 | 36.06 | 56.7 | - | 8.66 | 6.3 |
| Laurel (*Machilus thunbergii*) | 46.24 | 10.19 | 6.63 | 42.12 | 36.08 | 217.4 | - | 33.20 | 24.0 |
| Glossy privet (*Ligustrum lucidum*) | 29.32 | 10.55 | 12.98 | 24.50 | 18.40 | 42.8 | - | 6.54 | 4.7 |
| Hibiscus (*Hibiscus spp.*) | 22.83 | 13.74 | 12.64 | 12.97 | 14.67 | 29.4 | - | 4.48 | 3.2 |
| Japanese spindletree (*Euonymus japonicus*) | 32.36 | 12.59 | 7.84 | 28.07 | 24.61 | 63.7 | - | 9.73 | 7.0 |
| All browses | 39.10 | 10.28 | 7.93 | 37.51 | 31.19 | 491.7 |  | 75.1 | 54.2 |
| Apple | 14.62 | 3.87 | 2.03 | 9.92 | 8.28 | 23.5 | 98.3 | 3.58 | 2.6 |
| Carrot | 7.69 | 8.76 | 9.11 | 13.87 | 15.40 | 2.4 | 25.1 | 0.37 | 0.3 |
| Green bean | 5.44 | 12.95 | 23.88 | 24.65 | 24.28 | 3.9 | 41.4 | 0.59 | 0.4 |
| Broccoli | 10.81 | 9.16 | 32.11 | 14.27 | 13.61 | 15.7 | 98.3 | 2.40 | 1.7 |
| Asparagus | 4.52 | 12.40 | 29.97 | 26.26 | 19.75 | 4.4 | 95.9 | 0.67 | 0.5 |
| Cucumber | 2.54 | 16.92 | 21.39 | 18.75 | 19.49 | 3.0 | 97.3 | 0.46 | 0.3 |
| All fruits/vegetables | 8.48 | 7.78 | 16.31 | 14.33 | 12.95 | 52.8 | 79.5 | 8.1 | 5.8 |
| Soy bean | 39.95 | 5.91 | 45.44 | 17.73 | 17.25 | 16.3 | 99.7 | 2.48 | 0.3 |
| Peanuts | 95.65 | 3.69 | 26.01 | 44.24 | 38.20 | 44.5 | 63.9 | 6.80 | 4.9 |
| Primate L/S biscuit banana (Mazuri)*4 | 35.55 | 8.22 | 20.31 | 33.18 | 21.91 | 49.5 | 98.8 | 7.56 | 5.5 |
| Others | 48.69 | 6.05 | 26.32 | 35.37 | 27.80 | 110.3 | 79.0 | 16.8 | 12.2 |
| Ingested diet | 31.08 | 9.36 | 11.71 | 35.28 | 29.15 | 654.8 | - | 100.0 | 72.2 |

DM, dry matter; ash, crude ash; CP, crude protein; aNDFom, neutral detergent fibre assayed with a heat stable amylase exclude residual ash;
ADFom, acid detergent fibre excluded residual ash; DMI, dry matter intake; BW, body mass (kg)
\*1 Intake/offered of each feed (DM-basis). Browses were weighed as the whole branch, including leaves and twigs, but nutrient contents were analysed only for leaves. Thus, we could not calculate offered DM amounts of browse.
\*2 DM intake of each feed/total DM intake.
\*3 BW of the proboscis monkey was 18.9 kg.at the beginning of the second week of the experiment.
\*4 The pelleted feed was soaked in water before feeding

**S5 Table.**
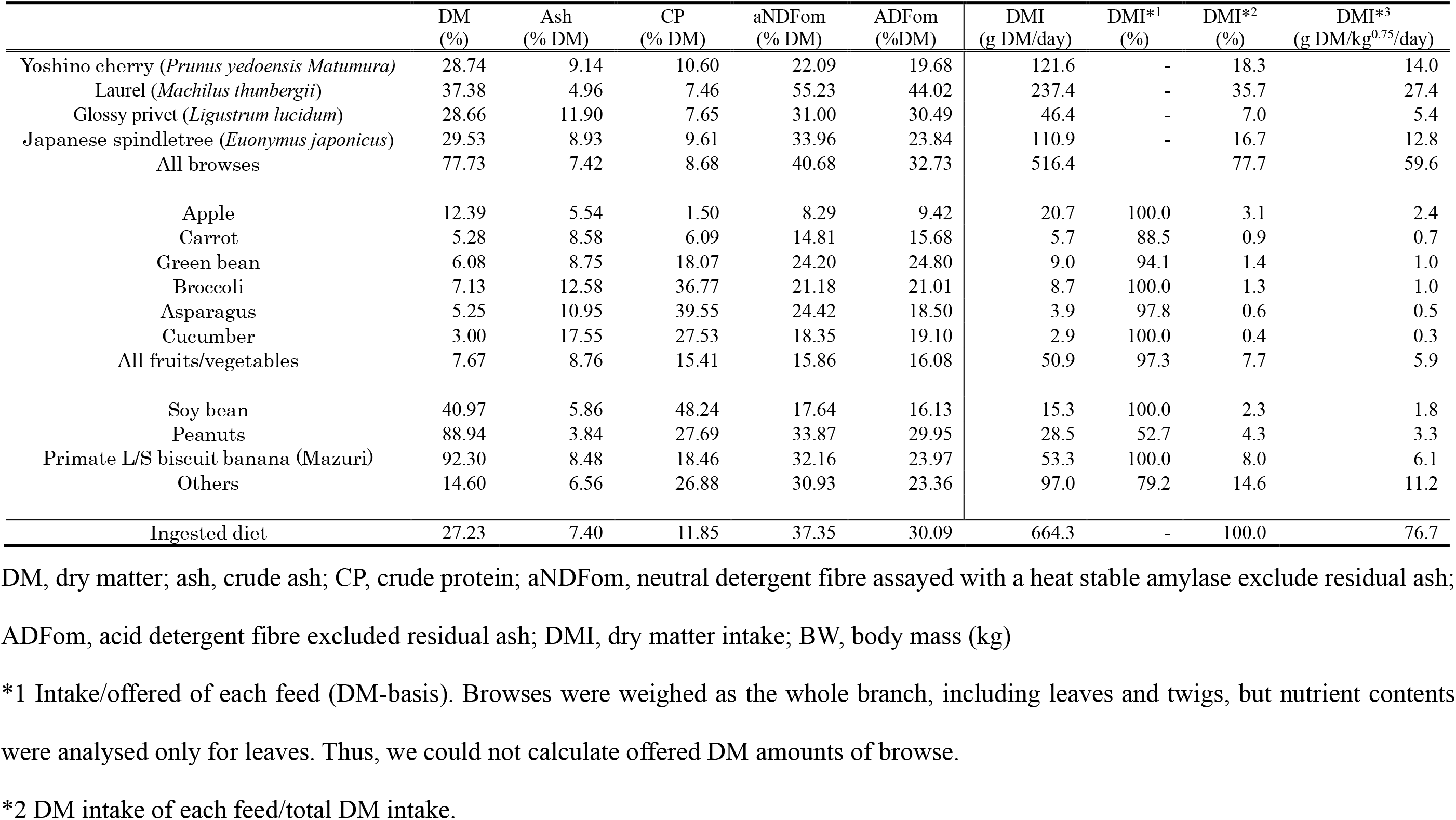
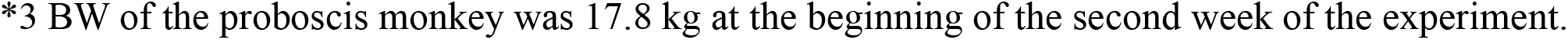
Nutrient contents in the diet and DM intake of an adult male of *Nasalis larvatus* in June 2018 (summer)

|  | DM<br>(%) | Ash<br>(% DM) | CP<br>(% DM) | aNDFom<br>(% DM) | ADFom<br>(%DM) | DMI<br>(g DM/day) | DMI*<sup>1</sup><br>(%) | DMI*<sup>2</sup><br>(%) | DMI*<sup>3</sup><br>(g DM/kg<sup>0.75</sup>/day) |
| --- | --- | --- | --- | --- | --- | --- | --- | --- | --- |
| Yoshino cherry (*Prunus yedoensis* Matumura) | 28.74 | 9.14 | 10.60 | 22.09 | 19.68 | 121.6 | - | 18.3 | 14.0 |
| Laurel (*Machilus thunbergii*) | 37.38 | 4.96 | 7.46 | 55.23 | 44.02 | 237.4 | - | 35.7 | 27.4 |
| Glossy privet (*Ligustrum lucidum*) | 28.66 | 11.90 | 7.65 | 31.00 | 30.49 | 46.4 | - | 7.0 | 5.4 |
| Japanese spindletree (*Euonymus japonicus*) | 29.53 | 8.93 | 9.61 | 33.96 | 23.84 | 110.9 | - | 16.7 | 12.8 |
| All browses | 77.73 | 7.42 | 8.68 | 40.68 | 32.73 | 516.4 |  | 77.7 | 59.6 |
| Apple | 12.39 | 5.54 | 1.50 | 8.29 | 9.42 | 20.7 | 100.0 | 3.1 | 2.4 |
| Carrot | 5.28 | 8.58 | 6.09 | 14.81 | 15.68 | 5.7 | 88.5 | 0.9 | 0.7 |
| Green bean | 6.08 | 8.75 | 18.07 | 24.20 | 24.80 | 9.0 | 94.1 | 1.4 | 1.0 |
| Broccoli | 7.13 | 12.58 | 36.77 | 21.18 | 21.01 | 8.7 | 100.0 | 1.3 | 1.0 |
| Asparagus | 5.25 | 10.95 | 39.55 | 24.42 | 18.50 | 3.9 | 97.8 | 0.6 | 0.5 |
| Cucumber | 3.00 | 17.55 | 27.53 | 18.35 | 19.10 | 2.9 | 100.0 | 0.4 | 0.3 |
| All fruits/vegetables | 7.67 | 8.76 | 15.41 | 15.86 | 16.08 | 50.9 | 97.3 | 7.7 | 5.9 |
| Soy bean | 40.97 | 5.86 | 48.24 | 17.64 | 16.13 | 15.3 | 100.0 | 2.3 | 1.8 |
| Peanuts | 88.94 | 3.84 | 27.69 | 33.87 | 29.95 | 28.5 | 52.7 | 4.3 | 3.3 |
| Primate L/S biscuit banana (Mazuri) | 92.30 | 8.48 | 18.46 | 32.16 | 23.97 | 53.3 | 100.0 | 8.0 | 6.1 |
| Others | 14.60 | 6.56 | 26.88 | 30.93 | 23.36 | 97.0 | 79.2 | 14.6 | 11.2 |
| Ingested diet | 27.23 | 7.40 | 11.85 | 37.35 | 30.09 | 664.3 | - | 100.0 | 76.7 |

DM, dry matter; ash, crude ash; CP, crude protein; aNDFom, neutral detergent fibre assayed with a heat stable amylase exclude residual ash;
ADFom, acid detergent fibre excluded residual ash; DMI, dry matter intake; BW, body mass (kg)
\*1 Intake/offered of each feed (DM-basis). Browses were weighed as the whole branch, including leaves and twigs, but nutrient contents were analysed only for leaves. Thus, we could not calculate offered DM amounts of browse.
\*2 DM intake of each feed/total DM intake.
\*3 BW of the proboscis monkey was 17.8 kg at the beginning of the second week of the experiment.

**S6 Table.** Nutrient contents in the diet and DM intake of two adults *Trachypithecus cristatus* in August 2018 (summer)

|  | DM<br>(%) | Ash<br>(% DM) | CP<br>(% DM) | aNDFom<br>(% DM) | ADFom<br>(%DM) | DMI<br>(g DM/day) | DMI*<sup>1</sup><br>(%) | DMI*<sup>2</sup><br>(%) | DMI*<sup>3</sup><br>(g DM/kg<sup>0.75</sup>/day) |
| --- | --- | --- | --- | --- | --- | --- | --- | --- | --- |
| Bamboo-leaf oak (*Quercus myrsinifolia*) | 51.48 | 7.60 | 9.19 | 56.62 | 34.56 | 16.0 | - | 2.0 | 1.7 |
| Apple | 14.17 | 6.12 | 1.53 | 5.24 | 6.24 | 73.5 | 95.9 | 28.1 | 8.0 |
| Banana peel | 8.81 | 17.68 | 8.48 | 30.10 | 30.67 | 6.8 | 94.6 | 3.2 | 0.7 |
| Cabbage | 8.74 | 8.62 | 15.24 | 14.23 | 14.50 | 69.8 | 98.5 | 21.7 | 7.6 |
| All fruits/vegetables | 10.76 | 7.80 | 8.22 | 10.55 | 11.19 | 150.1 | 96.9 | 52.9 | 16.4 |
| Carrot | 10.69 | 7.59 | 7.23 | 10.34 | 11.27 | 6.3 | 97.0 | 2.0 | 0.7 |
| Sweet potato | 31.66 | 3.33 | 2.74 | 6.88 | 3.55 | 101.3 | 99.2 | 34.2 | 11.1 |
| Leaf-Eater Primate Diet –<br>Mini-Biscuit (Mazuri) | 91.40 | 7.90 | 22.92 | 27.40 | 18.90 | 18.3 | 100.0 | 6.9 | 2.0 |
| Others | 31.54 | 4.21 | 5.90 | 10.03 | 6.16 | 126.0 | 99.3 | 43.1 | 13.8 |
| Ingested diet | 16.00 | 6.24 | 7.27 | 12.85 | 10.30 | 292.1 | - | 100.0 | 31.9 |

DM, dry matter; ash, crude ash; CP, crude protein; aNDFom, neutral detergent fibre assayed with a heat stable amylase exclude residual ash; ADFom, acid detergent fibre excluded residual ash; DMI, dry matter intake; BW, body mass (kg)
\*1 Intake/offered of each feed (DM-basis). Browsers were weighed as the whole branch, including leaves and twigs, but nutrient contents were analysed only for leaves. Thus, we could not calculate offered DM amounts of browse.
\*2 DM intake of each feed/total DM intake.
\*3 BW of two silver lutungs were 7.6 and 7.6 kg at the beginning of the second week of the experiment.

**S7 Table.**
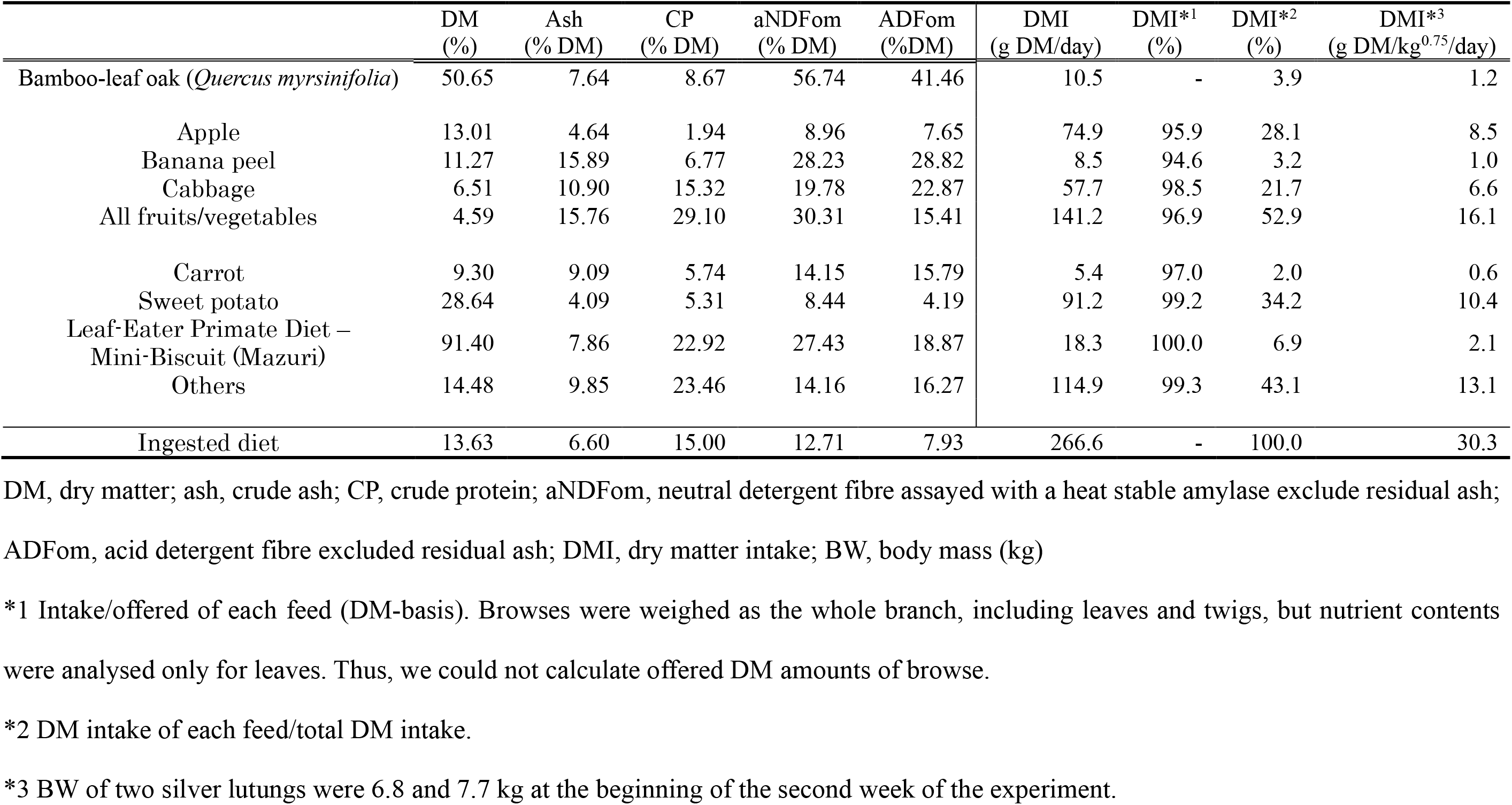
Nutrient contents in the diet and DM intake of two adults of *Trachypithecus cristatus* in February 2019 (winter)

